# Dotplotic: a lightweight visualization tool for BLAST+ alignments and genomic annotations

**DOI:** 10.1101/2025.05.12.653581

**Authors:** Hideyuki Miyazawa, Toshiyuki Oda

**Author notes:** **Corresponding Author:** Hideyuki Miyazawa.

## Abstract

With the development of sequencing technologies, chromosome-level genome assemblies have become increasingly common across various organisms, including non-model species. BLAST+ is one of the most widely used bioinformatics tools for computing sequence alignments, offering numerous optimizations for speed and scalability. Dot plots, which visualize the similarity between two sequences, are widely used in biological research. However, while many dot plot-generating programs exist, most rely on their own alignment algorithms, and it is uncommon to visualize external BLAST results directly.

Here, we present Dotplotic, a lightweight Perl program that generates dot plot-like visualizations based on BLAST output in tabular format. Dotplotic visualizes each alignment as a line connecting the start and end points of the query and subject sequences, with a gradient color indicating sequence identity. It allows users to overlay annotation data from external files onto the plot. Although command-line-based, Dotplotic requires only core Perl modules and is implemented as a single script, making it easy to install and run across platforms. The program supports standard input for both BLAST results and annotation files, enabling flexible visualization under various conditions, such as filtering specific alignments or displaying selected genomic features like genes or repeats.

Dotplotic is an efficient, portable, and easy-to-use visualization tool that enhances the exploration of sequence alignments and serves as a valuable resource for both bioinformatics and broader biological research.

## Background

Whole-genome sequencing has recently been conducted on a wide range of organisms because of advances in high-throughput technologies such as Oxford Nanopore and PacBio HiFi. Chromosome-level assemblies with high accuracy are becoming increasingly common—not only for the small genomes of viruses and prokaryotes but also for the large, repeat-rich genomes of vertebrates. Comparing assembled sequences, including non-coding regions such as introns and repeats, allows researchers to uncover genome diversity and evolution across species. For example, synteny patterns, such as gene order on chromosomes, are becoming increasingly important in phylogenetic studies (e.g., Schultz et al., 2023).

Dot plots are a widely used method for visualizing sequence comparisons and are helpful to researchers across many biological fields. For bioinformaticians, dot plots between assembled and reference sequences help assess the quality of sequencing and assembly. For evolutionary biologists, they reveal genomic rearrangements: large regions of similarity appear as diagonal lines, repeats generate multiple lines, inverted regions appear as lines in the opposite direction, and insertions or deletions appear as gaps.

In a basic dot plot, dots are placed where the two sequences share the same nucleotide or amino acid on a two-dimensional matrix. However, this simple method is unsuitable for large-scale sequence comparisons. Modern approaches instead calculate similarity between sequences and place dots in regions of high similarity. Several programs follow this approach, such as Dotter (Sonnhammer and Durbin, 1995), Gepard (Krumsiek et al., 2007), mummerplot (Marçais et al., 2018), FlexiDot (Seibt et al., 2018), and D-Genies (Cabanettes and Klopp, 2018). Because calculating sequence similarity is computationally intensive—particularly for searches involving many long sequences—these tools have improved their algorithms to reduce processing time and memory usage.

The Basic Local Alignment Search Tool (BLAST) is one of the most commonly used algorithms for sequence similarity searches (Altschul et al., 1990; 1997). BLAST+, a suite of command-line tools developed by the National Center for Biotechnology Information (NCBI), is widely used by both biologists and bioinformaticians (Camacho et al., 2009). It includes options for parallelization and for reducing database size by masking repetitive regions, making it suitable for large-scale searches on servers. Nearly all DNA sequences generated by researchers are stored in databases maintained by NCBI, and BLAST searches against this vast repository can be run through the NCBI website, contributing greatly to advances in biological and medical research.

Many software tools support BLAST searches and some generate dot plots based on BLAST output, including Unipro UGENE (Okonechnikov et al., 2012), Geneious (Kearse et al., 2012), and NBLAST (Choi et al., 2022). However, dot plot visualization tools that directly use external BLAST results remain scarce. Tools like Dotter rely on their own search algorithms and are not optimized to process BLAST output. GUI-based software such as UGENE is user-friendly but not well-suited for large-scale comparisons involving numerous chromosome-level sequences. Dotplotter, a CLI-based tool that can read external BLAST results (Mohite et al., 2025), offers limited functionality—for instance, it does not display sequence identity in the plots.

Here, we present Dotplotic, a command-line visualization tool that reads external BLAST output and generates dot plots. Dotplotic displays each alignment as a line colored according to sequence identity and allows users to overlay annotation data from external files. Output is generated in Scalable Vector Graphics (SVG) format, which can be edited with software such as Inkscape and exported to Portable Document Format (PDF).

Dotplotic is implemented as a single Perl script using only core modules and is hosted on GitHub, making it lightweight and easy to run on a server. This tool enables biologists familiar with the command line to easily visualize large-scale sequence comparisons.

## Implementation

Dotplotic is implemented in the Perl programming language (version 5.6 or later) and requires only core modules List::Util, Getopt::Long, and Pod::Text, making it compatible with a wide range of command-line environments, including macOS, Linux, and WSL2.

### Input data

BLAST+ programs (blastn, blastx, blastp, tblastn, and tblastx) support various output formats specified via the ‘outfmt’ option. Dotplotic supports three of these formats: tabular (‘-outfmt 6’), tabular with comment lines (‘-outfmt 7’), and comma-separated values (‘-outfmt 10’). Dotplotic requires that the BLAST search output be formatted appropriately using the ‘outfmt’ option. Users can include various fields in the BLAST output, such as ‘qaccver’ as Query accession version, and ‘bitscore’ as Bit score. The following nine fields are required for Dotplotic processing: ‘qaccver’, ‘saccver’, ‘pident’, ‘bitscore’, ‘length’, ‘qstart’, ‘qend’, ‘sstart’, and ‘send’. Dotplotic also supports overlaying annotation data on the plot, supporting GFF, BED, or OUT from RepeatMasker formats.

Minimap2, a sequence alignment program designed for large sequence datasets (Li, 2018), can output results in SAM format. The accompanying Perl script, Minimapsam2blast6.pl, converts this SAM output into a BLAST-like tabular format for compatibility with Dotplotic.

### Drawing process

Dotplotic represents each alignment as a single line (Fig. 1). Using four values_—_ query start (‘qstart’), query end (‘qend’), subject start (‘sstart’), and subject end (‘send’)_—_the program draws a line connecting the points (‘sstart’, ‘qstart’) and (‘send’, ‘qend’). The percentage of identical matches (‘pident’) is converted into a hexadecimal (hex) color code. Three identity thresholds_—_maximum (Max), middle, and minimum (Min)_—_are defined along with their corresponding color codes. By default, these are set to 100% (red, ‘#FF0000’), 80% (orange, ‘#FFFF00’), and 60% (lime, ‘#00FF00’). Identity values between Min and Max are linearly mapped to intermediate colors. Values below Min or above Max are assigned the Min and Max colors, respectively. Rectangles representing annotation features are drawn when annotation data is available.

**Fig. 1.**
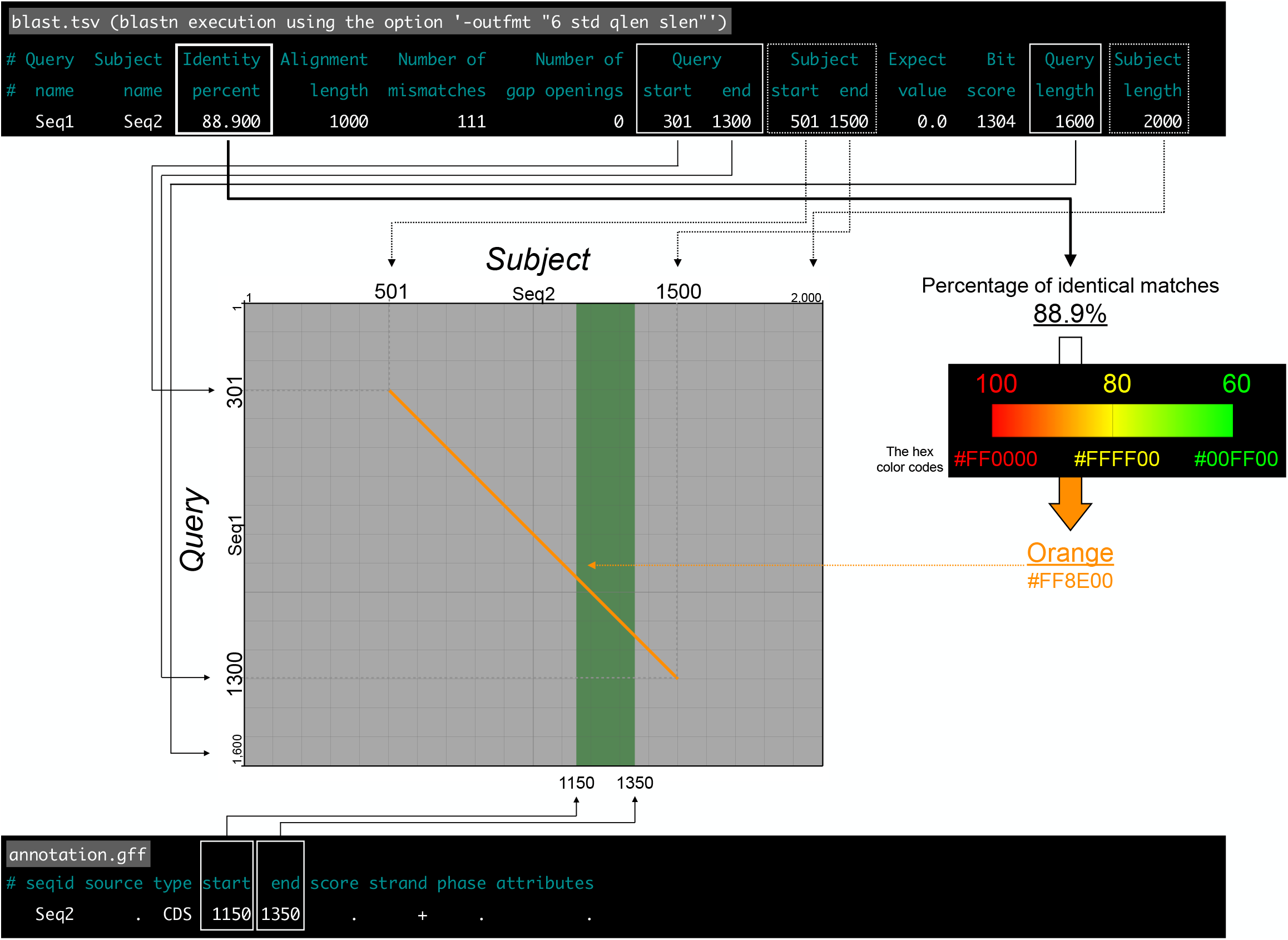
Visualization workflow of BLAST alignments and annotation data. Workflow for generating plots based on a BLAST alignment in tabular format and annotation data in GFF format. For each BLAST alignment, a line is drawn connecting the start and end positions of the query and subject sequences, with a hexadecimal color code representing sequence identity. A rectangle is also drawn to represent features from the annotation data.

The Dotplotic workflow consists of four steps (Fig. 2). Given a BLAST output file, and optionally an annotation file, Dotplotic reads the input, then calculates basic statistics, including the number and total lengths of query and subject sequences. If the sequence order is not specified by the user, Dotplotic performs a best-path search across all query-subject combinations to determine the query-subject relationships and their relative positions.

**Fig. 2.**
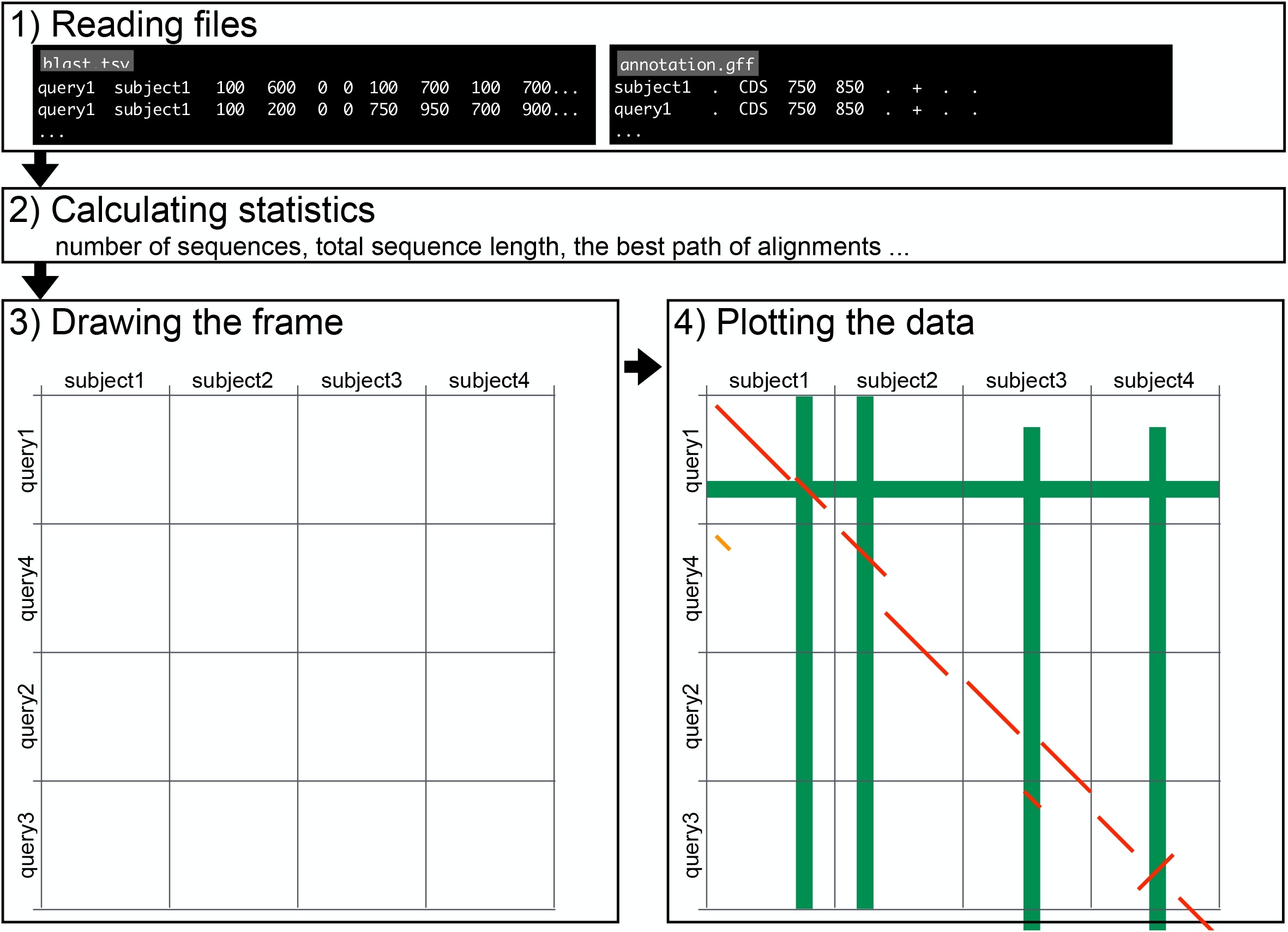
Overview of Dotplotic. Overview of Dotplotic. After reading the BLAST file (1), Dotplotic calculates the total sequence lengths and the number of query and subject sequences as well as similarities among them using a best-path search to determine the sequence order (2). Based on these results, frames are drawn, followed by the alignment lines and the annotation rectangles.

Subjects are sorted by name, and queries are ordered to form a continuous diagonal trend from the top left to the bottom right of the plot. For large BLAST files (e.g., genome-to-genome comparisons), best-path computation may consume significant memory and time. In such cases, users can reduce resource usage by manually specifying sequence order or by enabling the ‘light’ mode.

In the final plot, subjects are displayed along the horizontal axis and queries along the vertical axis, with annotation and alignment data rendered accordingly.

### Output format

Dotplotic produces output in SVG format, which can be opened in web browsers, such as Google Chrome and Mozilla Firefox. If the ‘click’ option is enabled, the SVG file embeds object information that can be viewed by clicking on graphical elements.

Vector graphic software such as Adobe Illustrator and Inkscape can open SVG files and export them to other formats. When converted to PDF, alignment lines and annotation rectangles remain sharp even when zoomed in. Although Dotplotic offers layout flexibility, the display may become cluttered if the BLAST input file includes too many sequences. To address this, the accompanying Perl script EditDotplotic.pl can be used to adjust visual elements such as font size or remove sequence length labels.

## Results

### Case study

Here, we demonstrate the visualization of BLAST alignments involving three yeast genomes using Dotplotic. The lager-brewing yeast *Saccharomyces pastorianus* is an interspecific hybrid of *S. cerevisiae* and *S. eubayanus*, and the genome of one of its strains, CBS 1483, has been sequenced at the chromosomal level (Salazar et al., 2019). We obtained nuclear genome sequence datasets for the three yeast species from GenBank: 31 chromosomes (total length: ca. 23.0 Mbp) for CBS 1483 (*S. pastorianus*, GenBank assembly accession: GCA_011022315.1), 16 chromosomes (total length: 11.6 Mbp) for *S. cerevisiae* (GCF_000146045.2), and 16 chromosomes (total length: 12.1 Mbp) for *S. eubayanus* (GCF_001298625.1). Additionally, we downloaded the genome annotation file of *S. cerevisiae* (GCF_000146045.2_R64_genomic.gff) from GenBank.

A BLAST search was conducted using CBS 1483 as the query and the other two species as subjects, with output in tabular format including query and subject lengths (using ‘-outfmt “6 std qlen slen”‘). Alignments shorter than 1 kbp were filtered out. Dotplotic was subsequently executed to visualize the BLAST alignments (Fig. 3). All commands used for the BLAST search and for running Dotplotic to generate Figures 3, 4, and 5 are listed in Supplementary Table 1. EditDotplotic.pl was also used to modify the sequence names in the Dotplotic outputs.

**Fig. 3.**
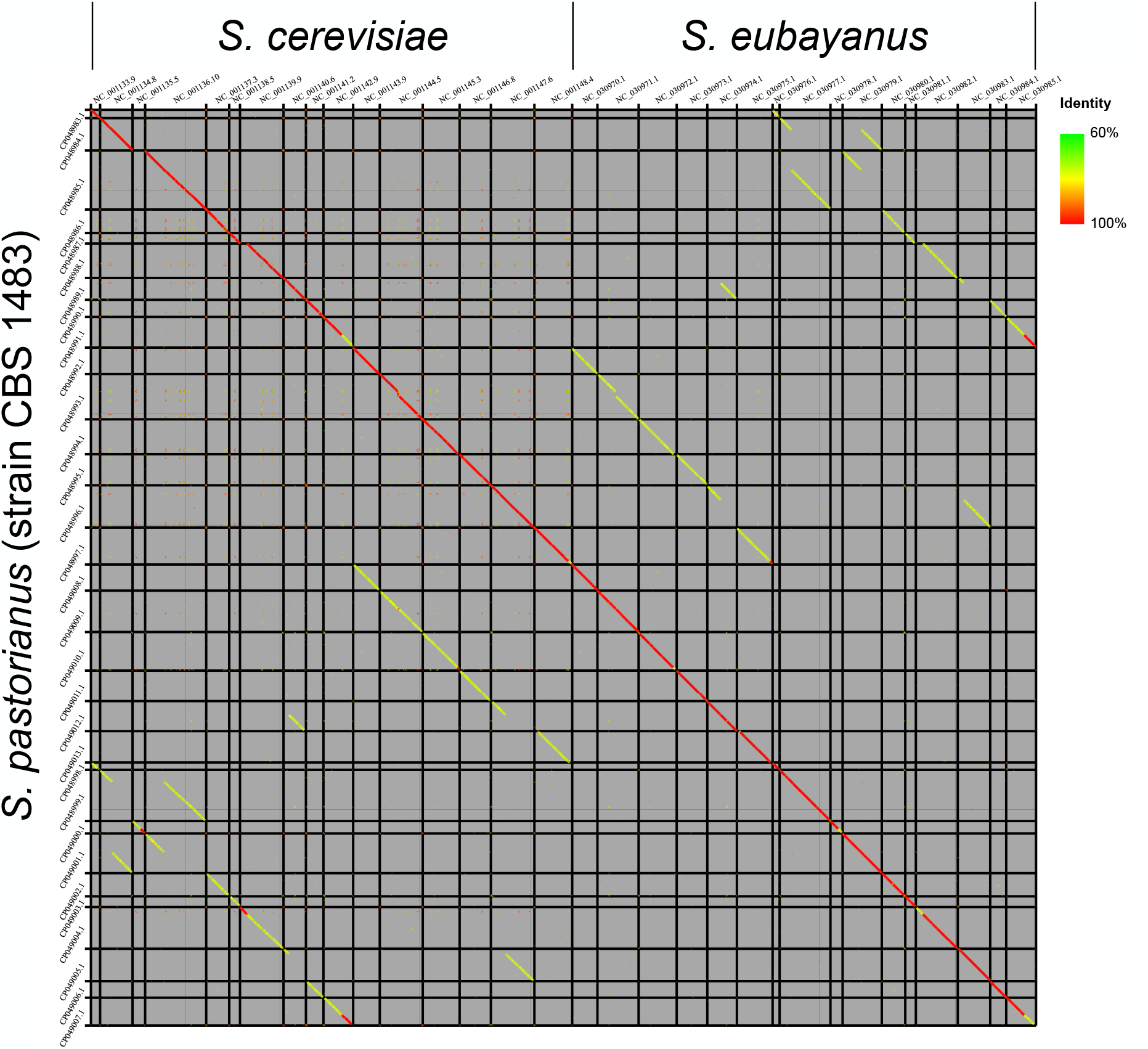
Genome comparison of yeasts using Dotplotic. The genome comparison of yeast genomes using Dotplotic.

**Fig. 4.**
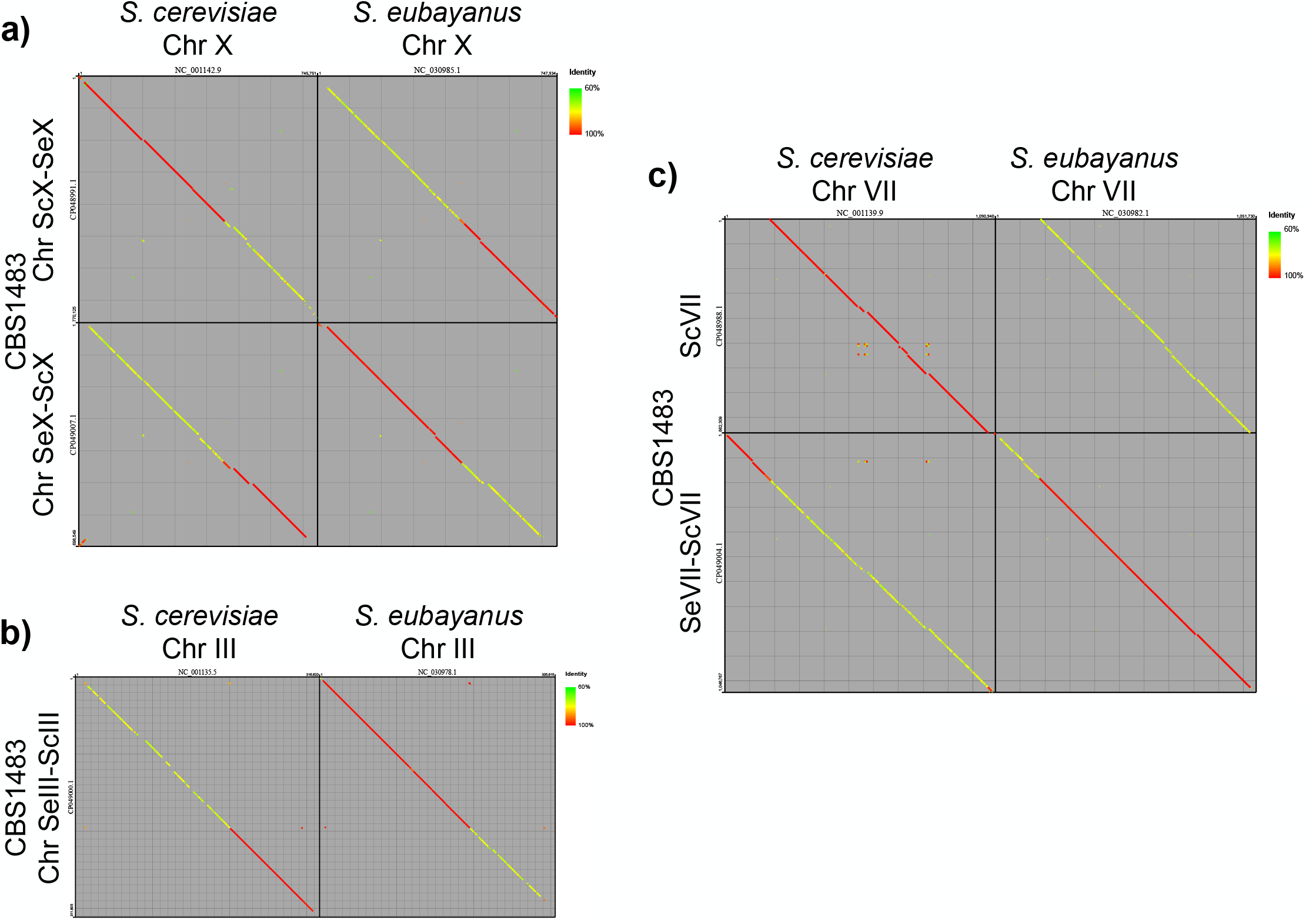
Chromosome comparison of yeasts using Dotplotic. Comparisons of yeast chromosomes using Dotplotic. Chromosomes are abbreviated as “Chr”.

**Fig. 5.**
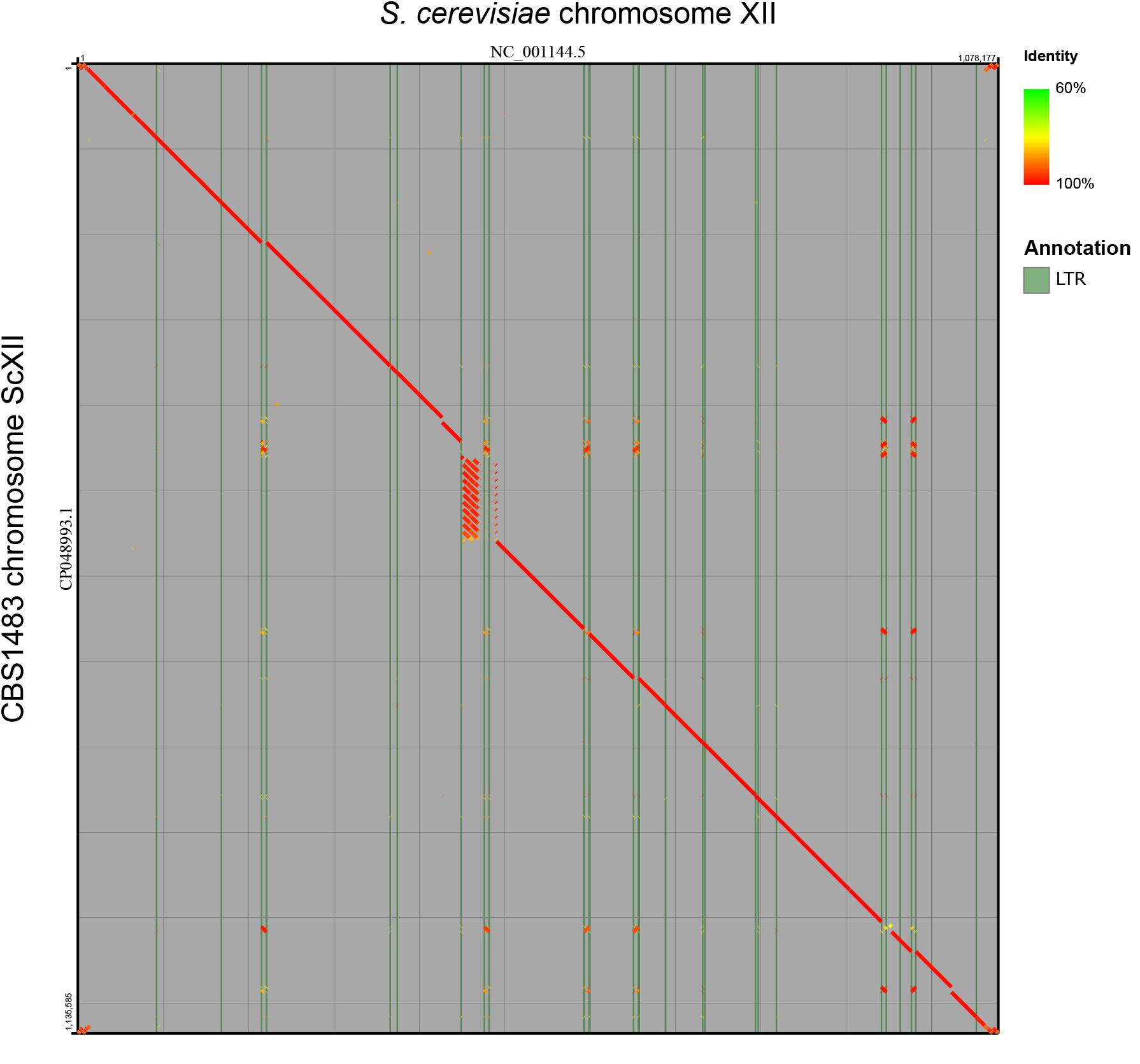
Chromosome comparison of yeasts with annotation data using Dotplotic.

Genomic rearrangements have been reported in several chromosomes (Salazar et al., 2019), and we visualized these events using Dotplotic (Fig. 4). Homologous recombination between chromosome X of *S. cerevisiae* and *S. eubayanus* was clearly observed in CBS 1483 (Fig. 4a). In chromosome SeIII-ScIII of CBS 1483, approximately two-thirds of the first half showed high similarity to chromosome III of *S. eubayanus*, while about one-third of the second half matched chromosome III of *S. cerevisiae*, suggesting a homologous recombination event between the two species (Fig. 4b). However, the truncated regions, approximately two-thirds of the first half of chromosome III in *S. cerevisiae* and about one-third of the second half of chromosome III in *S. eubayanus*, were not present in the CBS 1483 assembly (Fig. 3). While chromosomes ScVII and SeVII-ScVII in CBS 1483 correspond to chromosome VII in *S. cerevisiae* and *S. eubayanus*, respectively, approximately 100 kbp from the first half of *S. cerevisiae* chromosome VII align with that of SeVII-ScVII in CBS 1483, suggesting a translocation from *S. cerevisiae* to *S. eubayanus* (Fig. 4c). Although chromosome ScXII in CBS 1483 was generally similar to chromosome XII in *S. cerevisiae*, several repeated regions were observed (Fig. 5). The repetitive alignments in the central region suggest tandem repeat insertions. Additionally, all interspersed alignments were flanked by long terminal repeats (LTRs) on both the 5′ and 3′ ends, suggesting the proliferation of these regions along with LTR expansion.

## Discussion

Cabanettes and Klopp (2018) classified dot plot-generating tools into two generations. First-generation tools run only on Unix systems and produce static graphics without interactive features. Second-generation tools are platform-independent, user-friendly, and provide interactive interfaces, such as options for changing sequence orientation. The current version of Dotplotic runs solely on the command-line interface, which may not be user-friendly for many biologists. Although object information in the output can be viewed by enabling the ‘click’ option, the graphic itself is not interactive, indicating that Dotplotic belongs to the first-generation category. Furthermore, Dotplotic is solely a visualization tool and does not perform alignment searches independently.

Despite these limitations, we believe that Dotplotic offers three key advantages. First, it supports output from BLAST+, a widely used command-line alignment tool (Camacho et al., 2009). According to Google Scholar, BLAST+ was cited 2,890 times in 2024 alone, suggesting that it is routinely used by many bioinformaticians. By providing an intuitive visualization option for BLAST+ results, Dotplotic can significantly contribute to bioinformatics research.

Second, Dotplotic can easily overlay annotation data onto plots. The relationship between LTRs and repetitive regions is visualized in Fig. 5, which was generated with a single command (Table S1), demonstrating its ease of use. By modifying the annotation input, Dotplotic users can investigate the types and locations of genomic rearrangements.

Finally, Dotplotic is compatible with a wide range of platforms. Software with advanced features, such as high-throughput computing or multifunctional capabilities, is sometimes sensitive to changes in the development environment and may fail after updates to either the software or the underlying platform. In contrast, Dotplotic is a lightweight Perl program that depends only on core modules, making it likely to remain functional across diverse systems into the future.

## Conclusion

BLAST+ is a command-line–based alignment search tool and one of the most widely used programs in bioinformatics. However, most existing dot plot visualization tools rely on their own alignment algorithms, and visualizing results directly from BLAST+ output remains uncommon. To fill this gap, we present Dotplotic, a visualization tool specifically designed for BLAST+ output. Dotplotic is implemented as a single Perl script that depends only on core modules, making it lightweight, platform-independent, and easy to use.

Dotplotic can not only generate plots for genome comparisons but also overlay annotation data onto the plots, allowing users to visually compare genomic rearrangements with annotated features. This makes it a valuable tool for many bioinformaticians and biologists.

## Availability and requirements

Project name: Dotplotic

Project home page: https://github.com/HideyukiMiyazawa/Dotplotic

Operating system(s): MacOS, Linux.

Programming language: Perl

Other requirements: None

License:GPL-3.0 license

Any restrictions to use by non-academics: None

## Availability of data and materials

Dotplotic and other accompanying Perl programs, EditDotplotic.pl and Minimapsam2blast6.pl, are available on GitHub (https://github.com/HideyukiMiyazawa/Dotplotic)

## References

1. Altschul S. Gapped BLAST and PSI-BLAST: a new generation of protein database search programs. Nucleic Acids Research. 1997 Sep 1;25(17):3389–402.

2. Altschul SF, Gish W, Miller W, Myers EW, Lipman DJ. Basic local alignment search tool. Journal of Molecular Biology. 1990 Oct 5;215(3):403–10.

3. Cabanettes F, Klopp C. D-GENIES: dot plot large genomes in an interactive, efficient and simple way. PeerJ. 2018 Jun 4;6:e4958.

4. Choi BS, Choi SK, Kim NS, Choi IY. NBLAST: a graphical user interface-based two-way BLAST software with a dot plot viewer. Genomics Inform. 2022 Sep 30;20(3):e36.

5. Kearse M, Moir R, Wilson A, Stones-Havas S, Cheung M, Sturrock S, et al. Geneious Basic: An integrated and extendable desktop software platform for the organization and analysis of sequence data. Bioinformatics. 2012 Jun 15;28(12):1647–9.

6. Krumsiek J, Arnold R, Rattei T. Gepard: a rapid and sensitive tool for creating dotplots on genome scale. Bioinformatics. 2007 Apr 15;23(8):1026–8.

7. Marçais G, Delcher AL, Phillippy AM, Coston R, Salzberg SL, Zimin A. MUMmer4: A fast and versatile genome alignment system. Darling AE, editor. PLoS Comput Biol. 2018 Jan 26;14(1):e1005944.

8. Mohite OS, Jørgensen TS, Booth TJ, Charusanti P, Phaneuf PV, Weber T, et al. Pangenome mining of the Streptomyces genus redefines species’ biosynthetic potential. Genome Biol. 2025 Jan 14;26(1):9.

9. Okonechnikov K, Golosova O, Fursov M, the UGENE team. Unipro UGENE: a unified bioinformatics toolkit. Bioinformatics. 2012 Apr 15;28(8):1166–7.

10. Salazar AN, Gorter De Vries AR, Van Den Broek M, Brouwers N, De La Torre Cortès P, Kuijpers NGA, et al. Chromosome level assembly and comparative genome analysis confirm lager-brewing yeasts originated from a single hybridization. BMC Genomics. 2019 Dec;20(1):916.

11. Schultz DT, Haddock SHD, Bredeson JV, Green RE, Simakov O, Rokhsar DS. Ancient gene linkages support ctenophores as sister to other animals. Nature [Internet]. 2023 May 17; Available from: 10.1038/s41586-023-05936-6

12. Seibt KM, Schmidt T, Heitkam T. FlexiDot: highly customizable, ambiguity-aware dotplots for visual sequence analyses. Hancock J, editor. Bioinformatics. 2018 Oct 15;34(20):3575–7.

13. Shen W, Le S, Li Y, Hu F. SeqKit: A Cross-Platform and Ultrafast Toolkit for FASTA/Q File Manipulation. Zou Q, editor. PLoS ONE. 2016 Oct 5;11(10):e0163962.

14. Sonnhammer ELL, Durbin R. A dot-matrix program with dynamic threshold control suited for genomic DNA and protein sequence analysis. Gene. 1995 Dec;167(1–2):GC1–10.

