## Supplemental Table 1 for "Dotplotic: a lightweight visualization tool for BLAST+ alignments and genomic annotations"

### 1) preparation for the sequences

```
$ seqkit seq -m 100000 GCA_011022315.1_ASM1102231v1_genomic.fna > CBS1483.fa
$ seqkit seq -m 100000 GCF_000146045.2_R64_genomic.fna > TwoYeastGenomes.fa
$ seqkit seq -m 100000 GCF_001298625.1_SEUB3.0_genomic.fna >> TwoYeastGenomes.fa
```

**Note:** Only nuclear chromosomes were extracted from three yeast genome datasets with seqkit (Shen et al., 2016).

### 2) A BLAST search

```
$ blastn -task blastn -query CBS1483.fa -subject TwoYeastGenomes.fa \
    -outfmt "6 std qlen slen" | awk '1000 <= $4 {print}' > Blast.tsv
```

**Note:** The tabular format is specified using '-outfmt "6 std qlen slen", and short alignments less than 1 kbp are filtered using the 'awk' command.

### 3) Dotplotic

**Note:** To change the background color, the 'bg_color_set' option was set to '2' in the following commands.

```
$ Dotplotic --blast Blast.tsv --outfmt "6 std qlen slen" > Fig3.svg
```

```
$ Dotplotic --blast Blast.tsv --outfmt "6 std qlen slen" \
    --query CP048991.1,CP049007.1 --subject NC_001142.9,NC_030985.1 > Fig4a.svg
$ Dotplotic --blast Blast.tsv --outfmt "6 std qlen slen" \
    --query CP049000.1 --subject NC_001135.5,NC_030978.1 > Fig4b.svg
$ Dotplotic --blast Blast.tsv --outfmt "6 std qlen slen" \
    --query CP048988.1,CP049004.1 --subject NC_001139.9,NC_030982.1 > Fig4c.svg
```

**Note:** Sequences were specified using 'query' and 'subject' options.

```
$ awk '$3=="long_terminal_repeat" {print}' GCF_000146045.2_R64_genomic.gff | \
        Dotplotic --blast Blast.tsv --outfmt "6 std qlen slen" \
        --query CP048993.1 --subject NC_001144.5 \
        --annotation -:fmt=gff > Fig5.svg
```

**Note:** If the path to the BLAST result file is specified, Dotplotic can accept annotation data via standard input, with the file format specified (in this case, GFF). In this example, only annotation data for LTRs are displayed in the plot.
